# Integrative Analysis of Transcriptome Variation in Uterine Carcinosarcoma and Comparison to Sarcoma and Endometrial Carcinoma

**DOI:** 10.1101/012708

**Authors:** Natalie Davidson, Kjong-Van Lehmann, André Kahles, Alexander Perez, Gunnar Rätsch

## Abstract

Large-scale cancer genomics has made a huge impact onto cancer research. It has allowed the characterization of tumor types in an unprecedented depth. More recent studies target the joint analysis of multiple tumor types to gain insight into similarities and differences on a molecular level. Here we present an analysis of Uterine Carcinosarcoma. The histological similarities to sarcomas and carcinomas warrants an in-depth analysis to Uterine Endometrial Carcinoma as well as Sarcomas and we have used data from The Cancer Genome Atlas to understand transcriptome similarities and differences between these tumor types. We have performed a differential transcriptome analysis of Uterine Carinosarcoma to Uterine samples from GTEx to find genes with tumor specific splicing or expression patterns, which may not only be of interest for a deeper mechanistic understanding of the development and progression of Uterine Carcinosarcoma, but may also be potential tumor markers. Similarities and differences to Sarcomas and Endometrial Carcinomas present new opportunities for the development of new and targeted drug therapies. Finally we have also studied genetic determinants of gene expression and splicing changes and identified germline variants that explain expression and splicing differences between individuals. This analysis demonstrates the opportunities of integrative comparative analysis between multiple tumor types.

## 1. Introduction

Detailed molecular analysis of various tumor types has become feasible with the advent of large scale projects like The Cancer Genome Atlas (TCGA)^1^ providing an unprecedented resource of a variety of data modalities, allowing the investigation and comparison of multiple tumor types. This offers huge opportunities to gain insight into various cancer types and subtypes by comparing molecular similarities and differences. Recent efforts include the PanCancer initiative of the TCGA consortium^2^ releasing numerous companion manuscripts providing new insights into the molecular mechanisms underlying tumor development. Within those studies, transcriptome analysis on the level of gene expression has become mostly standard. However, changes of gene expression are only one type of derived transcriptome information influencing transcriptome complexity. Another major aspect of transcriptome complexity arises from the existence of multiple isoforms which are encoded by a single gene controlled by various RNA-modifying processes. The process of generating such alternate isoforms (alternative splicing) of a gene is highly controlled and involves the inclusion or exclusion of pre-specified parts of a gene.^3,4^ To what extent alternative splicing affects tumor progression and development is under investigation^5,6^ but it has already been shown that dysregulation and defects of the splicing processes play a role in cancer progression.^7–9^ These discoveries open up new treatment options since natural compounds as well as antisense oligonucleotides are in fact able to target aberrant splicing.^10,11^

Here, we present the results of a comparative study of transcriptome changes between 54 samples of uterine carcinosarcoma (UCS) with sarcoma (SARC), endometrial carcinoma (UCEC) and GTEx data from uterine tissue samples. We analyze the similarities of uterine carcinosarcoma to similar tumor types and tissue matching normal samples with respect to gene expression changes as well as aberrant splicing patterns. Further, we try to find somatic and germline mutations which determine changes in gene expression as well as splicing using a linear mixed model. This integrative analysis allows us to elucidate the differences and similarities between these tumor types.

## 2. Methods

This section only provides a brief description of methods that are crucial to the manuscript. For detailed, reproducible descriptions see Supplemental Information.

### 2.1. Data Processing

We have retrieved 54 UCS RNA-seq samples from the Cancer Genomics Hub (CGHub) and realigned them using STAR.^12^ In addition, we also retrieved RNA sequence data for 50 Sarcoma and 50 Uterine Corpus Endometrial Carcinoma samples as well as 36 samples taken from uterine tissue of non-cancer donors collected within the GTEx project. We processed all samples in a uniform manner. Based on the GENCODE annotation (version 19), we generated gene expression counts using a custom Python script and collected splice event quantifications with SplAdder.^13^

We also retrieved matching tumor/normal whole exome sequencing data for the UCS samples 70 and used the HaplotypeCaller^14^ as well as MuTect^15^ to identify germline and somatic sequence variants, respectively.

### 2.2. Principal Components Analysis

The Principal Components Analysis is based on the log transformed and library sized normalized expression quantification of all samples.

### 2.3. Tumor Specific Expression Analysis

UCS specific expression was assessed by first performing a gene quantification, then removing all genes whose expression is below the 40th quantile. Normalization of the remaining gene expression values was done using DESeq’s library size normalization method.^16^ After normalization, DESeq was used to call differentially expressed genes to compare a single UCS sample against all of the normal GTEx samples. All *p*-values were then FDR corrected using the Bonferroni method. Once a comparison was done for each tumor sample, a binomial test was used to test which genes were differentially expressed across the tumor samples. The binomial test compared the amount of samples in which each gene was found to be differential against a background acceptance rate for a random gene. After the differentially observed splicing events were found, GOrilla was used to do a functional enrichment analysis.^17^ To compare against UCEC and SARC samples, heatmaps were created to visually quantify patterns in gene expression.

### 2.4. Tumor Specific Splicing Analysis

UCS specific splicing was done by first using SplAdder^13^ to quantify alternative 3’ and alternative 5’ splice sites, intron retention, and exon skip events in the GTEx and UCS samples. The rDiff^18^ toolbox was then used to compare a single UCS sample against all of the normal samples. Once a comparison was done for each tumor sample, a binomial test was used to determine which splicing events were significant across all of the samples. The binomial test tested the amount of samples in which each splicing event was found to be differential against a background acceptance rate for a random splicing event. After the differentially observed splicing events were found, GOrilla was used to do a functional enrichment analysis.^17^

### 2.5. Expression and Splicing Quantitative Trait Analysis

Library size normalized expression quantification as well as splicing quantification measured in percent spliced in (PSI), both standard measures, are transformed via an inverse normal transform to ensure that deviations from the distributional assumption are not going to cause excess detection of false discoveries. Ties are resolved by adding random noise and an additive genetic model is used to encode variants. To account for population structure and other confounding factors we applied a linear mixed model.^19^ We used PANAMA^20^ to infer possible unknown confounders in addition to population structure.

## 3. Results

### 3.1. Molecular Transcriptome Signature of Uterine Carcinosarcoma Matches Endometrial Carcinoma

There has been a long standing discussion on the similarities of uterine carcinosarcoma in comparison to sarcoma and endometrial carcinoma, particularly since uterine carcinosarcoma exhibits features of both tumor types.^21–24^ Previously, uterine carcinosarcoma was classified as a uterine sarcoma due to histological features and aggressive behavior, but was recently reclassified as an endometrial carcinoma.^23^ Similarities between uterine carcinoma and uterine carcinosarcoma have been found in the level of p53 and p27 expression, lymph nodal involvement in malignancies, and risk factors.^21,24,25^

In order to understand the transcriptomic relationship between these tumor types we have analyzed the TCGA RNA-seq data. Figure 1 shows the samples projected on the first three principal components summarizing the relationship of these samples based on gene expression data. We observe that uterine carcinosarcoma samples seem to exhibit more similarity with endometrial carcinoma than with sarcoma samples on the first two principal components. This is consistent with the observation that uterine carcinosarcoma is more efficiently treated with therapies targeting endometrial carcinoma.^24^ Thus we conclude that the molecular signature based on transcriptomic data supports the idea that Uterine Carcinosarcoma show more similar patterns to Carcinomas than to Sarcomas.

**Fig. 1.**
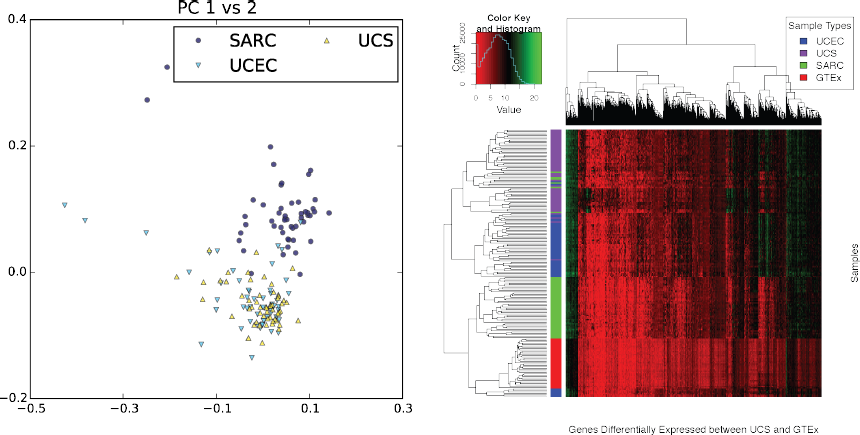
Left: First two principal components across thress cancer types based on expression data. Right: Hierachical clustering based on gene expression of about 1,700 genes.

### 3.2. Differential Gene Expression

There are a total of 2,308 differentially expressed genes between UCS and GTEx samples. Through the use of GOrilla, we found that there are several functional categories that are characteristic of most cancers such as regulation of cell cycle and cell division. More interestingly, we also found a few categories that seem to pertain specifically to UCS. These categories are: tissue morphogenesis (*q* < 3.7 × 10^−7^, where *q*-value has been corrected after Benjamini and Hochberg^26^), mesenchyme development (*q* < 8.04 × 10^−4^), and epithelial development (*q* < 1.49 × 10^−5^). We see in Figure 2, right panel, that when we cluster the expression of epithelium related genes, UCS and UCEC cluster together. Furthermore, there are a set of 11 genes that seem to visually distinguish UCEC and UCS from GTEx and SARC which are: GSTA1, PTHLH, NEUROG3, HYDIN, GSTA2, TBX1, GATA6, CNN3, DMBT1, ACADVL, DHRS9. These genes were not found to be alternatively spliced, nor known cancer genes. When we cluster the expression of sarcoma related genes, such as genes related to skeletal morphogenesis, shown in Figure 2, left panel, we do not observe very strong clustering between any one cancer type. This further suggests that genes related to the epithelium and thus carcinomas, act more similarly between UCS and UCEC, while sarcoma related genes do not show a similar pattern between the cancer types. Similarly to the heatmap related to the epithelium genes, we find 5 genes in the sarcoma genes that seem to visually distinguish UCS from all other samples: PLEKHA1, ANKRD11, NKX3-2, PDGFRB, IDUA. These genes are somewhat related to skeletal system morphogenesis, organ morphogenesis, and anatomical structure morphogenesis.

**Fig. 2.**
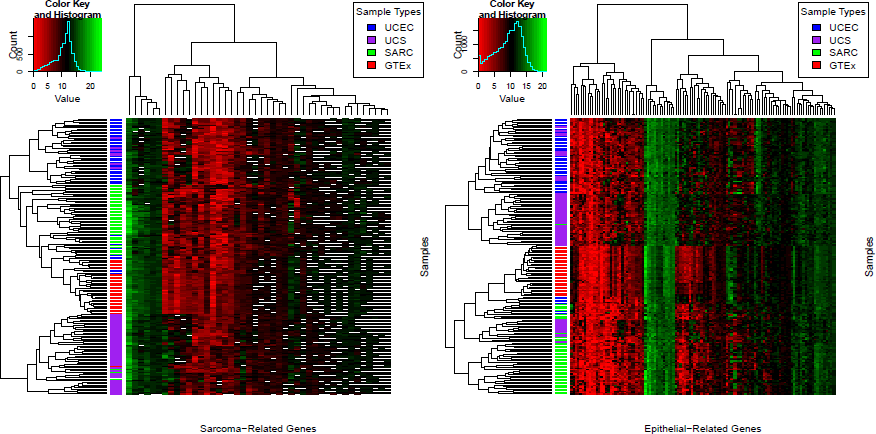
Clustering of cancer types of mesenchymal (left) and epithelial (right) genes

### 3.3. Differential Splicing

We find a total of 99 differentially spliced events between UCS and GTEx samples. Table 1 shows how many events were found for each type of splicing event. Of the 99 splicing events, we find 4 alternatively spliced genes that are associated with cancer: SRSF2, NACA, KMT2C, and TRIM27. Using GOrilla on a list of all the genes with splicing events ordered by the number of samples in which an event occured, we found several RNA-splicing related categories. The splicing related categories are: poly(A) RNA binding (*q* < 1.21 × 10^−6^, mRNA processing (*q* < 3.59 × 10^−2^, RNA splicing (*q* < 3.94 × 10^−2^), and regulation of mRNA stability (*q* < 8.31 × 10^−3^). This indicates that global splicing changes occur in UCS.

**Table 1.** Summary of alternative splicing events found to be differential between tumor and normal samples.

| Event Name | Number of Significant Events Found |
| --- | --- |
| Intron Retention | 54 |
| Exon Skip | 10 |
| Alternate 3' | 18 |
| Alternate 5' | 17 |

### 3.4. Tumor-Specific Splicing

We found a small set of introns that showed strong expression support in a large fraction of UCS tumor samples and could not be observed in any of the normal GTEx samples. To exclude the possibility that this observation is confounded by genes lacking expression in normal samples and thus explaining the absent introns, we restricted the analysis to genes that showed a higher mean expression in normals than in tumor samples. Note, that a gene can be highly expressed in normal samples but still none of the transcript isoforms contains the intron observed in the tumor samples. Figure 3 shows a ranked list of tumor specific introns that cannot be found in the GENCODE annotation (version 19). Although several genes have been associated to cancer before, e.g. SEC31A as part of gene fusions in lymphomas or SEC14L as progression marker for prostate cancer, none show a strong functional connection to UCS. An enrichment analysis of GO terms connected to the non-annotated tumor specific introns resulted in no significantly enriched categories. However, when including tumor specific introns that can be found in the annotation, we find a weak functional enrichment of cell-adhesion.

**Fig. 3.**
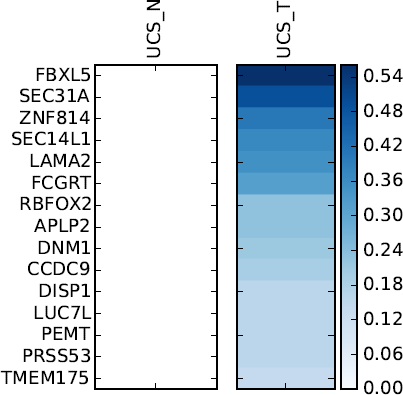
List of genes containing introns that are only observed in tumor but not in normal samples. Blue colorscale indicates fraction of tumor samples that show sufficient evidence of expression the tumor specific intron. UCS_T are the tumor samples and UCS_N are the normal samples taken from the GTEx project.

### 3.5. Genetic Determinants of Transcriptome Changes in Uterine Carcinosarcoma

While the previous sections described transcriptomic changes between tumor and normal data as well as a comparative analysis across multiple tumor types, this section explores potential genetic determinants of germline and somatic mutations in UCS. Due to the limited sample sizes available, we have restricted ourselves to study the effect of cis-associations only, in order to avoid excess false positive rates and control for the otherwise increased multiple hypothesis testing. Table 2 summarizes the amount of somatic as well as germline variants which are associated with expression and splicing changes. Considering the small sample size which significantly impacts the statistical power of this analysis, and the lack of whole genome data, we find a considerable amount of determinants of gene expression. We were unable to identify any somatic variants which are associated with either splicing changes or expression changes. Nevertheless, many of the germline alterations affecting splicing are associated with regulatory proteins (e.g., TCEA2,CHD3, EEF1D, RPL28, ZBTB8A). No enrichment of somatic variants could be found in these alternate splicing events. However, some alternate sequences have a length which is not divisble by three and thus point towards potential frameshift events.

**Table 2.** This table shows the number of cis-QTLs found across the various transcriptome changes, after multiple testing correction.

| Transcriptome Change | # QTLs | # Genes |
| --- | --- | --- |
| Exon Skip | 8 | 3 |
| Alternate 3' | 2 | 6 |
| Alternate 5' | 6 | 2 |
| Expression | 126 | 30 |

## 4. Discussion

In conclusion, we have done an extensive analysis of transcriptomic features on three tumor types. Our analysis improves our understanding of the relationship between UCS, UCEC and SARC on a molecular level and complements current knowledge which so far has been mostly based on histological analysis.^23^ Expression patterns confirm the similarities between UCS and UCEC and consistent with that observation we find that a clustering of samples based on epithelium related genes supports the similarity between UCS and UCEC, while mesanchyme related genes do not support any clustering between tumor types. Unfortunately the sample sizes are limited and thus an in-depth analysis of tumor subtypes was not possible and we were unable to study the characteristics in homologous and heterologous subtypes in UCS. An in-depth analysis of histological features and staging, in addition to transcriptomic changes could provide prognostic value that could be of interest to the community.

We identify differentially expressed genes and differentially spliced genes and provide a comparitive analysis to UCEC, SARC and GTEx normal samples. Interestingly we observe that alternatively spliced genes are enriched for splicing factors. We are confident that a better understanding of UCS in the light of UCEC and SARC, will lead to new insights and potential new drug targets. Nevertheless, larger sample sizes and tissue matched normal samples would contribute towards further insight. Here we have made use of transcriptomic data from GTEx, in order to do this differential analysis. Not only does the heterogeneity of the tumor sample pose challenges in this analysis, but also the differences between the data generation of the TCGA project and the GTEx project are potentially confounding our results. Not only has data been processed in different ways, but also TCGA data is extracted from live tissue, while GTEx data has been extracted post mortem. It is promising that despite confounding factors in our data, the trasciptomic variation we find reflect the similarities and differences between UCS, UCEC and SARC found previously using only histological, structural, and focused gene comparisons.

## Acknowledgements

We are grateful to the GTEx and TCGA consortia to provide the data to make this work possible. AK was funded through a fellowship of the Lucille Castori Center for Microbes, Inflammation and Cancer. Support was provided by the Tri-Institutional Training Program in Computational Biology and Medicine (via NIH training grant 1T32GM083937)

## References

1. J. Kaiser, Science 310, p. 1751 (2005).

2. J. N. Weinstein, E. A. Collisson, G. B. Mills, K. R. M. Shaw, B. A. Ozenberger, K. Ellrott, I. Shmulevich, C. Sander, J. M. Stuart, C. G. A. R. Network et al., Nature genetics 45, 1113 (2013).

3. M. Green, Annual Review of Genetics 20, 671(1986).

4. A. R. Kornblihtt, I. E. Schor, M. Alló, G. Dujardin, E. Petrillo and M. J. Muñoz, Nature Reviews Molecular Cell Biology 14, 153 (March 2013).

5. A. Valletti, O. Palumbo, M. D’Antonio, M. Gigante, M. Carella, E. Picardi, A. M. D’Erchia, R. Elena, G. Pesole and others, Cancer Epidemiology Biomarkers & Prevention 21, IA06 (2012).

6. D. Ramsköld, S. Luo, Y.-C. Wang, R. Li, Q. Deng, O. R. Faridani, G. A. Daniels, I. Khrebtukova, J. F. Loring, L. C. Laurent and others, Nature biotechnology 30, 777 (2012).

7. J. Tazi, N. Bakkour and S. Stamm, Biochimica et Biophysica Acta 1792, 14 (January 2009).

8. A. Srebrow and A. R. Kornblihtt, Journal of Cell Science 119, 2635 (2006).

9. V. Quesada, L. Conde, N. Villamor, G. R. Ordóñez, P. Jares, L. Bassaganyas, A. J. Ramsay, S. Beà, M. Pinyol, A. Martínez-Trillos, M. López-Guerra, D. Colomer, A. Navarro, T. Baumann, M. Aymerich, M. Rozman, J. Delgado, E. Giné, J. M. Hernández, M. González-Díaz, D. a. Puente, G. Velasco, J. M. P. Freije, J. M. C. Tubío, R. Royo, J. L. Gelpí, M. Orozco, D. G. Pisano, J. Zamora, M. Vázquez, A. Valencia, H. Himmelbauer, M. Bayés, S. Heath, M. Gut, I. Gut, X. Estivill, A. López-Guillermo, X. S. Puente, E. Campo and C. López-Otín, Nature Genetics 44, 47 (January 2012).

10. S. Bonnal, L. Vigevani and J. Valcárcel, Nature Reviews Drug Discovery 11, 847 (November 2012).

11. B. Khoo and A. Krainer, Current Opinion in Molecular Therapeutics 11, 108 (2009).

12. A. Dobin, C. a. Davis, F. Schlesinger, J. Drenkow, C. Zaleski, S. Jha, P. Batut, M. Chaisson and T. R. Gingeras, Bioinformatics 29, 15 (January 2013).

13. X. Gan, O. Stegle, J. Behr, J. G. Steffen, P. Drewe, K. L. Hildebrand, R. Lyngsoe, S. J. Schultheiss, E. J. Osborne, V. T. Sreedharan, A. Kahles, R. Bohnert, G. Jean, P. Derwent, P. Kersey, E. J. Belfield, N. P. Harberd, E. Kemen, C. Toomajian, P. X. Kover, R. M. Clark, G. Rätsch and R. Mott, Nature 108, 10249 (August 2011).

14. M. A. DePristo, E. Banks, R. Poplin, K. V. Garimella, J. R. Maguire, C. Hartl, A. A. Philippakis, G. del Angel, M. A. Rivas, M. Hanna, A. McKenna, T. J. Fennell, A. M. Kernytsky, A. Y. Sivachenko, K. Cibulskis, S. B. Gabriel, D. Altshuler and M. J. Daly, Nature Genetics 43, 491 (2011).

15. K. Cibulskis, M. S. Lawrence, S. L. Carter, A. Sivachenko, D. Jaffe, C. Sougnez, S. Gabriel, M. Meyerson, E. S. Lander and G. Getz, Nature Biotechnology 31, 213 (2013).

16. S. Anders and W. Huber, Genome Biology 11, p. R106 (2010).

17. E. Eden, R. Navon, I. Steinfeld, D. Lipson and Z. Yakhini, BMC Bioinformatics 10, p. 48 (2009).

18. P. Drewe, O. Stegle, L. Hartmann, A. Kahles, R. Bohnert, A. Wachter, K. Borgwardt and G. Rätsch, Nucleic acids research 41, 5189 (2013).

19. C. Lippert, F. Casale, B. Rakitsch and O. Stegle, bioRxiv , 0 (2014).

20. N. Fusi, O. Stegle and N. Lawrence, Nature Precedings , 1 (2011).

21. A. Gadducci, S. Cosio, A. Romanini and A. R. Genazzani, Critical reviews in oncology/hematology 65, 129 (2008).

22. F. Amant, I. Cadron, L. Fuso, P. Berteloot, E. de Jonge, G. Jacomen, J. Van Robaeys, P. Neven, P. Moerman and I. Vergote, Gynecologic oncology 98, 274 (2005).

23. E. D’Angelo and J. Prat, Gynecologic oncology 116, 131 (2010).

24. W. McCluggage, Journal of clinical pathology 55, 321 (2002).

25. A. Abargel, I. Avinoach, V. Kravtsov, M. Boaz, M. Glezerman and J. Menczer, International Journal of Gynecological Cancer 14, 354 (2004).

26. Y. Benjamini and Y. Hochberg, Journal of the Royal Statistical Society. Series B (Methodological) 51, 289 (1995).

